# *In vitro* discovery of a therapeutic lead for HFMD from a library screen of rocaglates/aglains

**DOI:** 10.1101/2024.09.30.615979

**Authors:** Adrian Oo, Angel Borge, Regina Ching Hua Lee, Cyrill Kafi Salim, Wenyu Wang, Michael Ricca, Deborah Yuhui Fong, Sylvie Alonso, Lauren E. Brown, John A. Porco, Justin Jang Hann Chu

## Abstract

The lack of an effective antiviral treatment for enteroviruses, including the human enterovirus A71 (EV-A71), has resulted in an immense global healthcare burden associated with hand-foot-and-mouth disease (HFMD). Rocaglates and aglains belong to a family of compounds produced by *Aglaia* genus plants. Since the initial discovery of rocaglates in 1982, various rocaglates and aglains have been synthesized and extensively studied as anticancer and antiviral agents. Here, we report our studies towards the discovery of a novel aglain derivative as an EV-A71 inhibitor and work to decipher its antiviral effect. From an immunofluorescence-based phenotypic screen of a library of 296 rocaglate and aglain derivatives, we identified a lead aglain derivative which effectively suppressed EV-A71 replication by 2.3 log fold at a non-cytotoxic concentration. Further validation revealed inhibition of EV-A71 across multiple cell types and a pan-enterovirus inhibitory spectrum against other enteroviruses. Subsequent mechanistic investigation revealed interference with EV-A71 intracellular post-entry events including viral RNA transcription and translation. Findings from this study have established a strong foundation for development of aglain scaffolds as much needed antiviral agents for HFMD, paving the way for future medicinal chemistry optimization and *in vivo* studies.

## 1. Introduction

Hand, foot and mouth disease (HFMD) is an infectious viral disease commonly affecting infants and children which results from infection by viruses belonging to the *Picornaviridae* family^1^. Enterovirus A71 (EV-A71), Coxsackievirus A16 (CV-A16), and CV-A6 are the primary etiological agents of HFMD, in addition to the less prevalent CV-A4, CV-A9, CV-B1, and CV-B5^2,3^. Although HFMD has been typically self-limiting with onsets of fever, ulcers, and rashes which generally resolve within a week, potentially fatal neurological and respiratory complications are major health concerns of the disease^1,4^. Unlike vector-borne diseases such as Dengue fever, HFMD outbreaks are not geographically restricted, and are easily transmitted *via* contaminated droplets, faeces, or surfaces of objects. HFMD outbreaks have been reported worldwide annually with seasonal peaks and sporadic cases in between, resulting in significant paediatric hospitalization and an associated healthcare burden^5,6^. To date, clinically available EV-A71 vaccines are solely approved for usage in China and provide restricted prophylactic coverage, limited to the C4 genotype^7^. This limitation, combined with the current lack of an approved antiviral agent for HFMD, underscore the crucial need for development of novel and effective therapeutic agents for this disease.

In this study, we examined rocaglates (flavaglines) and aglain scaffolds^8^ against EV-A71. Rocaglates are cyclopenta[*b*]benzofurans produced by plants from the genus *Aglaia* (**Figure 1A**) and include the antiproliferative natural products silvestrol (**1**) and rocaglamide A (RocA, **2**). Rocaglates are effective in blocking viral replication in cell culture through potent clamping of the mammalian DEAD-box helicase eIF4A onto its substrate RNA, leading to inhibition of protein translation^9^. This mechanism has been leveraged across multiple therapeutic areas ranging from oncology to infectious diseases^10^, including EV-A71^11^. A number of synthetic rocaglates including the hydroxamate **CR-1-31b** (**3**) synthesized by the Porco laboratory^12^ (**Figure 1B**) have also been shown to have potent antiviral activity^13,14^. Furthermore, we have recently reported inhibition of SARS-CoV-2 replication by synthetic ‘amidino rocaglates’ (ADRs) such as **4**^15^. There have been several literature reports of rocaglates with promising antiviral activities against various major human viruses, including a recent report on halogenated rocaglates with inhibitory effects against hepatitis E and SARS-CoV-2 viruses^16^.

**Figure 1.**
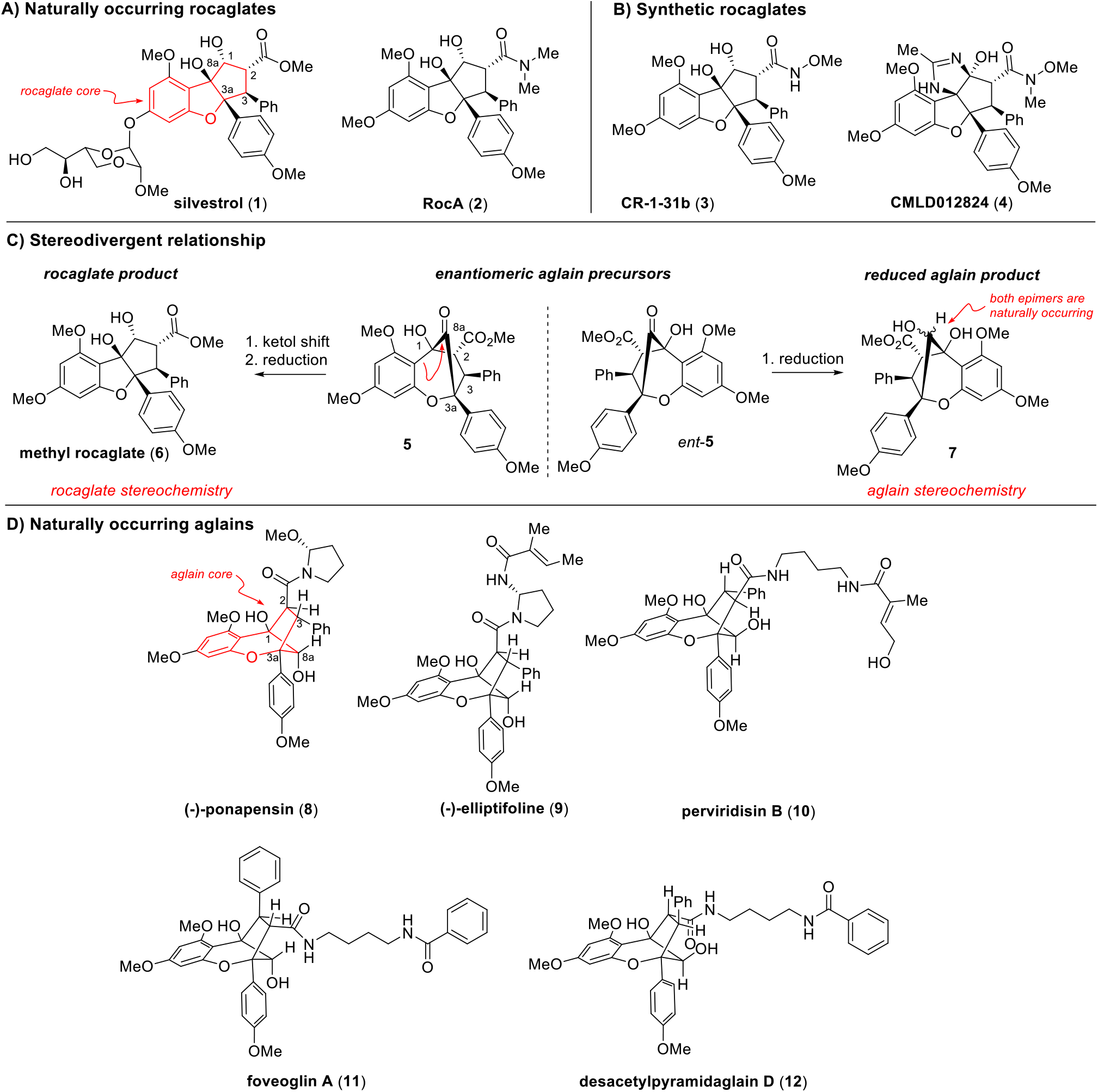
Chemical structures of (A) naturally occurring rocaglates (**1** and **2**), and (B) synthetic rocaglates (**3** and **4**); (C) Proposed biosynthetic transformation of aglain ketone precursors **5***/ent*-**5** to rocaglates such as **6** (atoms and rings are labelled according to the typical convention for rocaglates). Enantioselective total synthesis and bioactivity profiles have so far established that bioactive aglains and bioactive rocaglates are derived from aglain precursors that are enantiomeric to one another. (D) Chemical structures of naturally occurring aglains **8-12**.

A second chemical class also isolated from *Aglaia* genus plants, aglains, are cyclopenta[*bc*]benzopyrans which have been proposed to serve as biosynthetic precursors to rocaglates^17^ (**Figure 1C**). Compared to rocaglates, the activity spectrum of aglains has been more sparsely characterized. Interestingly, the enantioselective total syntheses of both rocaglate and aglain natural products by the Porco laboratory revealed the possibility of stereochemical divergence for the scaffolds^18^ (**Figure 1C**), suggestive that bioactive rocaglates and bioactive aglains are likely derived from separate enantiomers of a common chiral, racemic aglain precursor (±)-**5**. To-date, all known bioactive rocaglates possess absolute stereochemistry that is transitively analogous to aglain precursor **5**. In contrast, two bioactive aglains have been shown by synthesis to possess absolute stereochemistry matching that of the enantiomeric aglain precursor *ent*-**5**. For example, the naturally-occurring aglain (-)-ponapensin (**8, Figure 1D**), isolated from *A. ponapensin* and shown by synthesis to be derived from *ent*-**5**, exhibits potent NF-κB inhibition (IC50 = 60 nM)^19^. Additionally, (-)-elliptifoline (**9**), isolated from *A. elliptifolia* and also synthetically derived from *ent*-**5**, exhibits weak cytotoxic activity against A549 and P-388 cell lines (ED50s of 18.9 µg/mL and 3.4 µg/mL, respectively)^20^. Additional bioactive aglains include perviridisin B (**10**)^21^, isolated from *A. perviridis* with cytotoxicity against HT-29 cells (ED50 = 0.46 µM) and weak NF-κB inhibition (2.4 µM), and foveoglin A (**11**), a cytotoxic aglain isolated from *A. foveolata* with 1.4-1.8 µM activity against lung, prostate, and breast cancer cell lines^22^. To date, there has only been a single report of an antiviral aglain natural product, desacetylpyramidaglain D (**12, Figure 1D**), which was found to be a modest inhibitor of herpes simplex virus (HSV)^23^. Based on the reported activities of both rocaglates and aglains, we evaluated a collection of 296 synthetic variants of these chemotypes from the Boston University Center for Molecular Discovery (BU-CMD) compound collection against EV-A71.

## 2. Experimental Section

### 2.1. Compounds and Reagents

A total of 264 rocaglate and 32 aglain derivatives were provided by the Boston University Center for Molecular Discovery (BU-CMD). Compounds were dissolved in dimethyl sulfoxide (DMSO) to a 10 mM concentration and were stored at -80 °C prior to use in assays. The materials and methods to produce chiral, racemic tested aglains **10, 11**, and **13**-**18** have been previously described^18,24-26^. All compounds carried forward to secondary assays were determined to be >95% pure by LC/MS with ELSD detection. Experimental details for the X-ray crystal structure of compound **13** are provided in the Supplementary Information.

### 2.2. Cells and Viruses

Cell lines used in this study include murine motor neuron cells (NSC-34; CELLutions Biosystems, CLU140), human neuroblastoma cells (SH-SY5Y; ATCC®CRL-2266™) and human muscle rhabdomyosarcoma cells (RD; ATCCCCL-136). All cell lines were maintained in Dulbecco’s Modified Eagle’s Medium (DMEM) in the presence of 10% heat-inactivated fetal calf serum (HI-FCS) and 2 g sodium hydrogen carbonate at 37 °C, with 5% CO2 (incubation condition referred to as “incubated” hereafter). Enteroviruses targeted in this study were EV-A71 strain 41 (Accession no. AF316321.2); CV-A6 (Accession No. KC866983.1); and CV-A16 (Accession No. U05876). All enteroviruses were propagated in RD cells with reduced serum DMEM. Virus stocks were kept in aliquots at -80 °C until needed for subsequent experiments.

### 2.3. Preliminary Screen

Each rocaglate/aglain derivative was subjected to preliminary evaluation of their respective cytotoxicity profiles and antiviral inhibitory potential against EV-A71, as published^27^. Cytotoxicity screening was conducted using the alamarBlue™ Cell Viability Reagent (Thermo Fisher Scientific, Waltham, MA, USA) while each compound’s antiviral potential was determined using the Operetta High-Content Imaging System (PerkinElmer, Waltham, MA, USA). The percentage infection of each treatment or control group and robustness of our preliminary screening assay based on Z-factor were evaluated as previously described^27^.

### 2.4. Cytotoxicity and Antiviral Validation

The complete cytotoxicity profiles of selected rocaglate/aglain derivatives against NSC-34 or SH-SY5Y cells were determined via the alamarBlue™ assay. Treated cells were incubated with the compounds for 12 h (NSC-34) or 24 h (SH-SY5Y). Negative and vehicle controls of this assay involved cells treated with fresh reduced serum DMEM and 0.1% DMSO, respectively. Cell viability in each treatment group was determined based on the intensity of colorimetric changes of each treatment group. For antiviral validation, cells were infected with EV-A71 (MOI 1) and incubated for 1 h. Following incubation, infected cells were rinsed with sterile 1X PBS to remove any unbound virus particles. Specific concentrations of respective compounds were added to the cells in triplicates and incubated for 12 h (NSC-34) or 24 h (SH-SY5Y). At the end of the incubation period, two cycles of freeze-thaws (-80 °C; 37 °C) were performed on the plates prior to collection of supernatants from each treatment or control group for virus titre measurement via plaque assay, as described^27^.

### 2.5. Temporal-based Antiviral Assays

The specific phase(s) of EV-A71 replication cycle during which our hit compound exerts its optimal inhibitory effects were evaluated with a series of time-dependent approaches: time-of-addition (TOA), time-of-removal (TOR), pre-treatment, co-treatment and entry-bypass assays, as published^27^. The assays were performed in SH-SY5Y cells in triplicates and resulting virus titres were quantified *via* plaque assays as previously described^27^.

### 2.6. Quantitative Reverse Transcription-Polymerase Chain Reaction (qRT-PCR)

Total RNA was extracted from each treatment and control group using the RNeasy Mini Kit (QIAGEN). The resulting RNA samples were subjected to reverse transcription (M-MLV Reverse Transcriptase, Promega) to obtain cDNA of either the positive-or negative-strand of EV-A71 RNA using the forward (5’-CCTCCGGCCCCTGAATGCGGCTAAT-3’) and reverse primers (5’-ATTGTCACCATAAGCAGCCA-3’), according to the manufacturer’s protocol. Respective resulting cDNA fragments were utilized for qPCR analyses involving the same primer sets using SYBR Green Quantitative RT-PCR kit (Sigma-Aldrich). Quantification of β-actin mRNA (forward: 5’-TCGGTGAGGATCTTCATGAGGTA-3’; reverse: 5’-TCACCCACACTGTGCCCATCTACGA-3’) was simultaneously performed as the endogenous control, against which the viral RNA strands were normalized. Reactions were carried out in the Applied Biosystems StepOnePlus qPCR system according to the manufacturer’s protocol. Data was analysed using the Livak method^28^ to obtain fold-changes of compound-treated samples relative to DMSO-treated samples.

### 2.7. SDS-PAGE and Immunoblotting

Cells from specific treatment and control groups were lysed with lysis buffer containing 1X Laemmli buffer. The resulting cell lysates were heated for 10 min at 95 °C and subjected to SDS-PAGE run in 10% acrylamide gels at 100 V for 2 h. The proteins were then transferred to a polyvinylidene fluoride membrane using the Trans-Blot Turbo system (Bio-Rad). Following a 20 min membrane blocking using 2% bovine serum albumin (BSA) dissolved in Tris-buffered saline-Tween 20 (TBST), specific viral protein and host loading controls were probed using the mouse antibodies targeting EV-A71 VP2 (MAB979) and host β-actin, respectively. After 1 h incubation with constant rocking at room temperature, the membranes were subjected to three rounds of washes using TBST prior to incubation at room temperature with horseradish peroxidase (HRP) conjugated goat anti-mouse IgG secondary antibody (Thermo Fisher Scientific) for 1 h. Following three rounds of membrane washing with TBST, specific protein bands were developed using the Immobilon Western Chemiluminescent HRP substrate (Merck Millipore) for 3 min. Band visualization and analyses were performed using the C-DiGit Chemiluminescence Western Blot Scanner and Image Studio™ Version 4.0 (LI-COR, Lincoln, NE, USA), respectively.

### 2.8. Nano-Luciferase Replicon Assays

Using the DharmaFECT 1 transfection reagent (Dharmacon), SH-SY5Y cells seeded in 96-well plates were transfected with either our established replication-competent or replication-defective EV-A71 replicon construct^27,29^ and incubated for 4 h. At the end of incubation, the transfected cells were washed with sterile 1X PBS and treated with 0.1% DMSO or respective concentrations of test compound or positive controls. Guanidine hydrochloride (GuHCl) was used as positive control for the replication-competent replicon, and cycloheximide (CHX) was used as positive control for the replication-defective replicon. After 24 h, luciferase signal detection was subsequently performed using the Nano-Glo kit (Promega, Madison, WI, USA) with a GLOMAX Multi-Detection System microplate reader (Promega, Madison, WI, USA).

### 2.9. Statistical Analyses

GraphPad Prism 9 (GraphPad Software, Inc.) was used for data analyses. One-way analysis of variance (ANOVA) followed by a Dunnett’s post hoc test were performed to determine the significance of each reading relative to its respective control in each experimental setup. Samples exhibiting statistical difference from the control exhibit *p*-values of < 0.05 (^*^), < 0.01 (^**^) and < 0.001 (^***^).

## 3. Results

### 3.1. High-Throughput Phenotypic Screen of EV-A71 Inhibitors

To identify potential EV-A71 antiviral molecules from a library of 296 rocaglates and aglains, a high-throughput immunofluorescence assay was performed as previously described^27^. Treatment-induced cell death was simultaneously evaluated *via* an alamarBlue™ assay to prevent false positive antiviral hits as viable cells are required for productive virus replication. Compounds were assessed for reduction in EV-A71 viral titre and host cell cytotoxicity at 10 µM. The robustness of our high throughput antiviral screening platform was validated *via* Z-factor assessment (Z-factor: 0.55) as described previously^27^. Our typical “hit” criteria in this screen is defined as compounds reducing EV-A71 viral titre by >2 logs in the absence of host cell cytotoxicity (>90% host cell viability). Only one compound, aglain **13**, met this threshold (**Figures 2A-B**). This compound, bearing a pentafluorophenyl (PFP) ring^30-32^, was prepared as part of a methodology study towards the synthesis of the isomeric aglain natural products perviridisin B (**10**)^24^ and foveoglin A (**11**). We have also fully confirmed the structure of **13** by X-ray crystal structure analysis (**Figure 3**).

**Figure 2.**
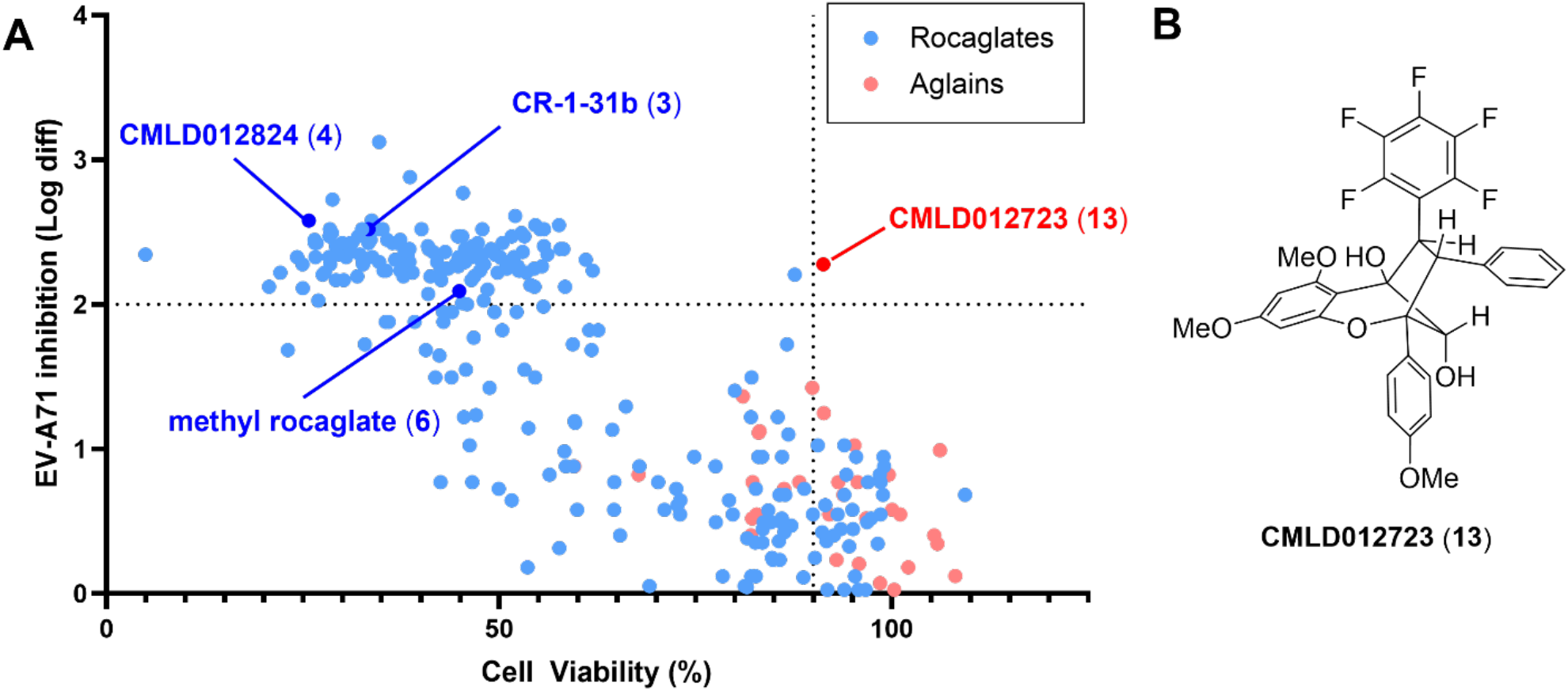
Summary of EV-A71 activity screen. For the cytotoxicity screen, NSC-34 cells were treated with 10 µM of each compound and incubated for 12 h prior to cell viability evaluation using the alamarBlue™ Cell Viability Reagent (Thermo Fisher Scientific, Waltham, MA, USA). Fluorescence intensity in each treatment and control group was measured at 570 nm excitation wavelength and 600 nm emission wavelength using the Infinite™ 200 series microplate reader (Tecan, Männedorf, Switzerland). For the antiviral screen, NSC-34 cells were infected with EV-A71 (MOI 1) and incubated for 1 h, followed by treatment with 10 µM of respective compounds for 12 h within a similar incubation environment. Methanol-fixed cells were probed with anti-EV-A71 VP2 1° antibodies (MAB979; Merck Millipore, Burlington, VT, USA) and anti-mouse FITC 2° antibodies (Merck Millipore), prior to being visualized using the Operetta High-Content Imaging System. Image analyses were performed with the Harmony High-Content Imaging and Analysis Software (PerkinElmer, Waltham, MA, USA). Resulting images were captured with DAPI and FITC fluorescence filters from a pre-determined central locus of each well and processed with the Cell Profiler Software which generates quantitative readings on the total number of cells as well as the infected cell population within each well via signals from the DAPI-stained nuclei and FITC-stained EV-A71 VP2, respectively. (A) Scatter plot of host cell viability (X-axis) and antiviral activity (Y-axis) with data points colored by scaffold type. All tested rocaglates with >2 log reduction were unacceptably toxic to host cells, including exemplar rocaglates **3, 4** and **6**. Only one aglain hit (**CMLD012723, 13**) met the established hit criteria of >2 log reduction in viral titre with >90% cell viability. (B) Chemical structure of aglain hit compound **13**.

**Figure 3.**
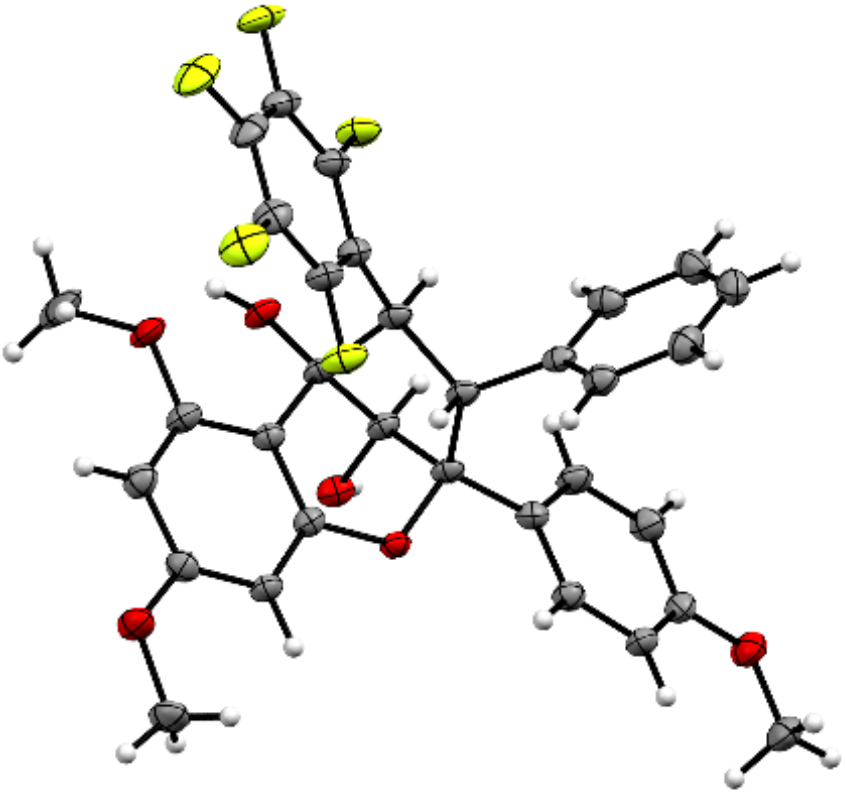
ORTEP drawing of the single crystal X-ray structure of aglain **13**. Thermal ellipsoids are shown at the 50% probability level. Diffraction data were collected on a Bruker D8 Venture, and the ORTEP was generated using Mercury.

Examining the activity profile of the remaining compounds screened, it was clear that a high level of host cell cytotoxicity was a limiting factor for the vast majority of rocaglates that showed appreciable antiviral activity (**Figure 2**, *blue*), including exemplar rocaglates **3, 4** and **6** (**Figures 1B-1C)**. We thus opted to focus on the aglains (**Figure 2**, *red*), which collectively showed significantly reduced cellular toxicity. By focusing the field to aglains and expanding our “hit” criteria to >1 log reduction in viral titre, we identified several additional congeners of interest (**Figures 4A-B**). For example, synthetic, racemic variants of the aglain natural products perviridisin B (**10**) and foveoglin A (**11**) reduced viral titre by 1.4 and 1.0 logs, respectively. Compound **14**, a des-fluorinated variant of **13** that is also oxidized at the 2° hydroxyl to the corresponding ketone, similarly showed a modest (∼1 log) reduction in viral titre. Lastly, we noted that compound **15**, an “*aza*-aglain”^26^ so named for the oxygen-to-nitrogen scaffold substitution producing a cyclopenta[*bc*]tetrahydroquinoline core, showed 1.2-log reduction in EV-A71 titre. Of this expanded field of five hits, we noted that four compounds (**10, 11, 13** and **14**), all bore an aromatic substituent at the “C2” position (see **Figure 1C** for atom numbering), albeit with different stereochemistry for **10**, which is diastereomeric to the other aglains at both C2 and C3. In contrast, *aza*-aglain **15** bore a carbomethoxy ester at the C2 position. In examining other *aza*-aglains tested in the screen, we noted that the closely related *aza*-aglain **16** (**Figures 4A-B**), synthesized from **15** by diastereoselective reduction of the bridge carbonyl to afford a 2° alcohol epimeric to **13**, also showed antiviral activity with excess cytotoxicity (81% cell viability at 10 µM). We also noted compound **17** with significant structural similarities to both **15** (C2 ester) and **13** (B-ring 4-methoxyphenyl substitution and similar 2° alcohol stereochemistry), which was found to be non-cytotoxic and showed no measurable antiviral activity.

**Figure 4.**
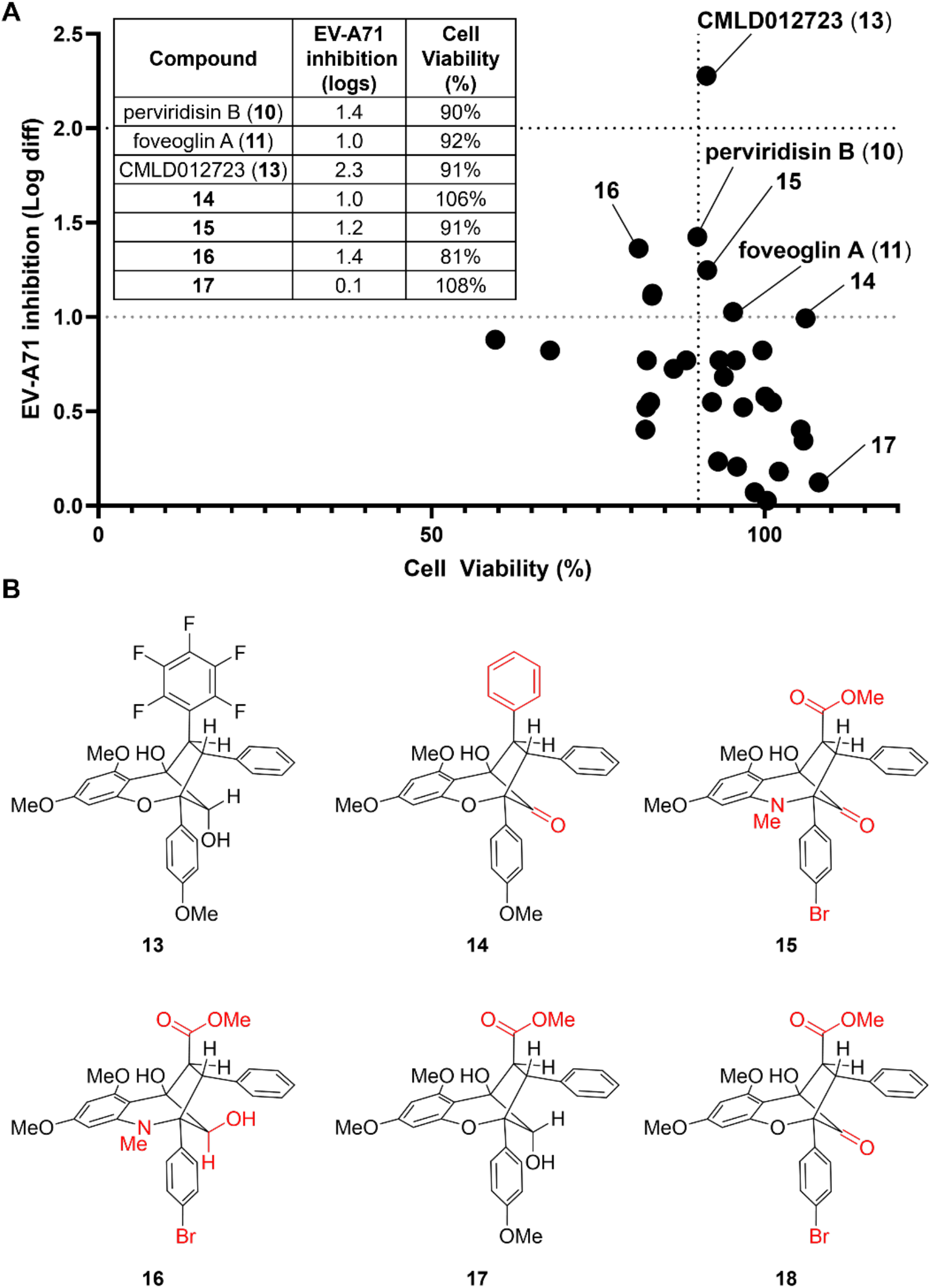
(A) Expanded hit criteria and near-neighbour identification for non-cytotoxic aglains reveals additional compounds of interest **10, 11** and **14-17**. (B) Chemical structures of compounds **13-18**, with key structural differences from screening hit **13** highlighted in red.

### 3.2. Validation of Hit Aglains

While a shortage of available material precluded our advancement of perviridisin B (**10**), we carried the remaining active cohort (**11, 14**, and **15**) as well as our top hit **13** forward to secondary plaque-based antiviral assays to validate their respective dose-dependent activity for both viral replication inhibition and host cell cytotoxicity. To better explore structure-activity relationships (SAR) governing antiviral action, we also targeted *aza*-aglain **16** and the published, unscreened compound **18**^18^, which was chosen as a “non-*aza*” variant of **15**. As illustrated in **Figure 5A**, the only aglain showing significant reduction in EV-A71 titre was compound **13** (EC50: 3.57 µM; **Figure S1**), which also showed host cell cytotoxicity that precluded antiviral testing at concentrations higher than 10 µM. Apart from compound **18**, which exhibited a significant degree of cytotoxicity, foveoglin A (**11**) as well as other tested compounds were non-toxic at all tested concentrations. However, these compounds were generally less potent than compound **13** as depicted by respective higher EC50 values. Hence, compound **13** was selected as a lead compound for further evaluation.

**Figure 5.**
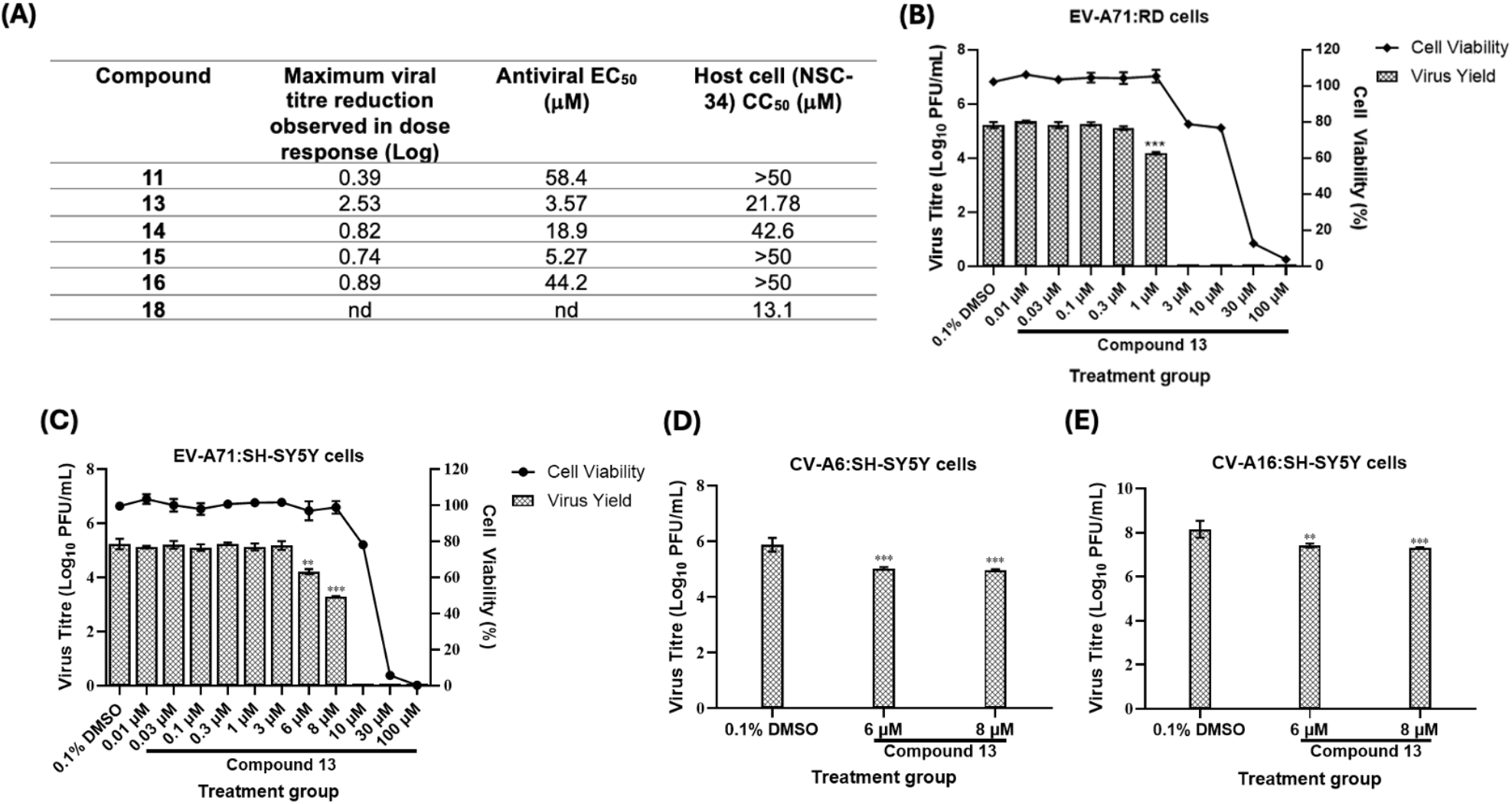
Validation of cytotoxicity and antiviral profiles of selected aglain derivatives. (A) NSC-34 cells were treated with various concentrations of respective compounds, in the presence or absence of EV-A71 (MOI 1) infection. Resulting cell viability and virus yield from respective experimental setups were determined via alamarBlue™ and plaque assays, respectively. EV-A71 inhibition potency and cytotoxicity of each compound was tabulated whereby EC50 and CC50 values were computed using GraphPad Prism 9 software. Dose-response evaluation of cytotoxicity and antiviral properties of compound **13** against EV-A71 in (B) RD or (C) SH-SY5Y cells. In SH-SY5Y cells, potential pan-enterovirus spectrum of compound **13** was investigated against (D) CV-A6 or (E) CV-A16. Every assay was performed in triplicates with error bars representing the standard deviation from the mean of triplicates. 0.1% DMSO served as the mock control group of respective assays.

As the initial screening and validation assays were performed in the murine-derived NSC-34 cells, we next sought to confirm compound **13**’s antiviral efficacy against EV-A71 in human cell lines to better understand its therapeutic potential for human infections. In order to exclude potential cell-specific bias of the observed antiviral activity, compound **13**, was evaluated for anti-EV-A71 activity in both the neuronal cell line SH-SY5Y, and the highly susceptible RD cells^33,34^. Compound **13** showed more favourable anti-EV-A71 inhibition in both cell lines; the virus replication was suppressed at an EC50 of 0.43 µM in RD cells (**Figure 5B**) in comparison with NSC-34 cells (EC50 = 3.57 µM; **Figure 5A**), whereas approximately 2 logs downregulation of virus titer was achieved at a lower non-cytotoxic concentration of the compound in SH-SY5Y cells (8 µM; **Figure 5C**, vs 15 µM in NSC-34 cells; **Figure S1**). Although a more promising selectivity index (SI = CC50/EC50) was computed for RD cells (SI = 25.98) than SH-SY5Y cells (SI = 2.02), subsequent downstream assays were performed in the neuronal cells given the severe clinical significance of neurologic complications associated with HFMD. The pan-enteroviral potential of compound **10** was also investigated by testing against CV-A6 and CV-A16, two other key pathogens causing HFMD. At the two highest non-cytotoxic concentrations (6 µM and 8 µM), compound **13** was observed to effectively reduce infectious CV-A6 **(Figure 5D)** and CV-A16 **(Figure 5E)** titers, suggesting that the compound may hold promise as a broad-spectrum anti-enterovirus inhibitor.

### 3.3. Antiviral Evaluation during Specific Stages of EV-A71 Replication Cycle

To gain further understanding on the underlying mechanism(s) involved in compound **13**’s antiviral activity against EV-A71, a series of temporal-based assays representing the different phases of the virus life cycle were performed. The time-window of EV-A71 life cycle during which compound **13** exerted its antiviral activity was first determined by TOA and TOR assays. This approach involved the addition (TOA) or removal (TOR) of compound **13** within the surrounding environment of infected cells at different time points post-infection until 18 h.p.i, at which point supernatants from each treatment group and controls were collected for viral titre quantification by plaque assays. From our TOA findings, EV-A71 yields were suppressed when **13** was added prior to 6 h.p.i. whereas continuous reduction in the viral load was observed when treatment was halted across every tested time point up to 18 h.p.i. **(Figure 6A)**. Subsequent pre-**(Figure 6B)** and co-treatment **(Figure 6C)** assays revealed ineffective inhibition of EV-A71 replication by compound **13** when infected cells were exposed to the compound prior to, or during, virus infection of host cells. Interestingly, EV-A71 was significantly downregulated when **13** was administered following transfection of its viral RNA into target cells as shown by our entry-bypass assay **(Figure 6D)**. From this experimental setup, we observed that 6 µM and 8 µM of compound **13** effectively suppressed virus yields by 0.88- and 2.27 logs, respectively.

**Figure 6.**
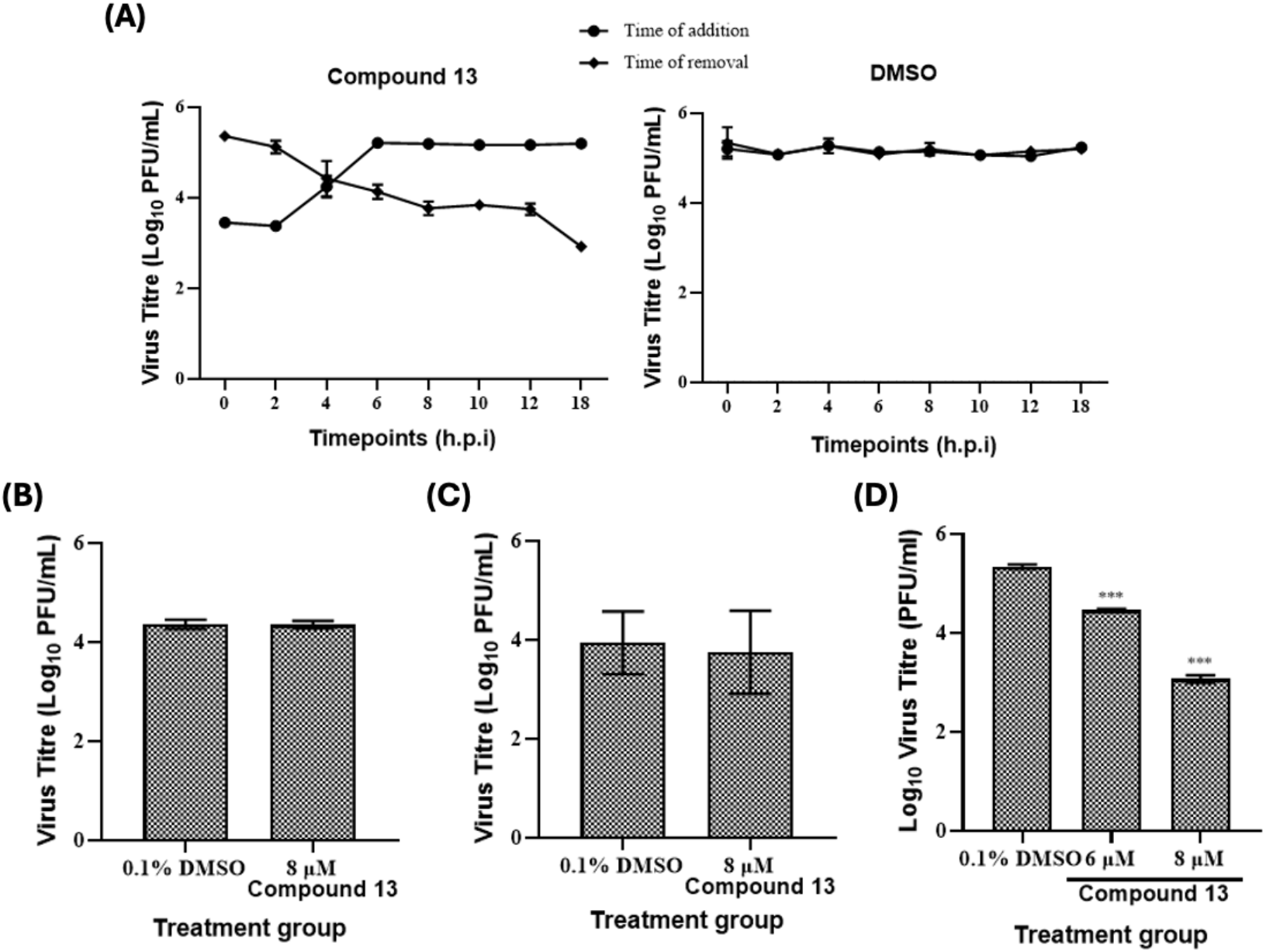
Temporal-based evaluation of compound **13**’s antiviral activity against EV-A71. (A) Time-of-addition (TOA) and time-of-removal (TOR) of compound **13** (8 μM) at specific time points across the EV-A71 replication cycle in SH-SY5Y cells. For TOA, virus-containing media was aspirated and replaced with compound **13** (8 μM) at 0, 2, 4, 6, 8, 10, 12 or 18 h post-infection (h.p.i.). For TOR, infected SH-SY5Y cells were first incubated with compound **13** (8 μM), and media was decanted and replaced with fresh media at similar time points as TOA. At 24 h.p.i., supernatant from each treatment and control group was collected for virus quantification via plaque assay. Separately, SH-SY5Y cells were infected with EV-A71 (MOI 1) and treated with specific concentrations of compound **13** according to previously published experimental schemes for (B) pre-treatment, (C) co-treatment and (D) entry-bypass assays^27^. At 24 h.p.i., supernatant was collected and used for virus yield measurement via plaque assay. Every assay was performed in triplicates with error bars representing standard deviation from the mean of triplicates. 0.1% DMSO served as the mock control group of respective assays.

### 3.4. Antiviral Evaluation against EV-A71 RNA replication and Protein Translation

To assess compound **13**’s modulatory effects on EV-A71 RNA synthesis and protein production, we employed qRT-PCR, immunoblotting and replicon-based approaches. Consistent with our plaque assay-quantified virus yields, significant decrease in both positive- and negative-sense RNA of EV-A71 were detected following treatment with 6 µM and 8 µM of aglain **13** at 12 and 24 h.p.i. **(Figure 7A)**. The effective reduction in viral RNA was subsequently translated into prominent downregulation of viral protein expression levels (> 50%) as evidenced by weaker band intensities of EV-A71 VP2 in cells treated with the compound in comparison with the vehicle control **(Figure 7B)**. Using our established replication-competent and replication-defective EV-A71 replicons^27^, we then observed that compound **13** significantly decreased the luciferase signals generated from both constructs **(Figure 7C)**. In fact, stronger inhibition of the replication-competent replicon was observed in the presence of 6 and 8 µM concentrations of compound **13** than that of the replication-defective construct.

**Figure 7.**
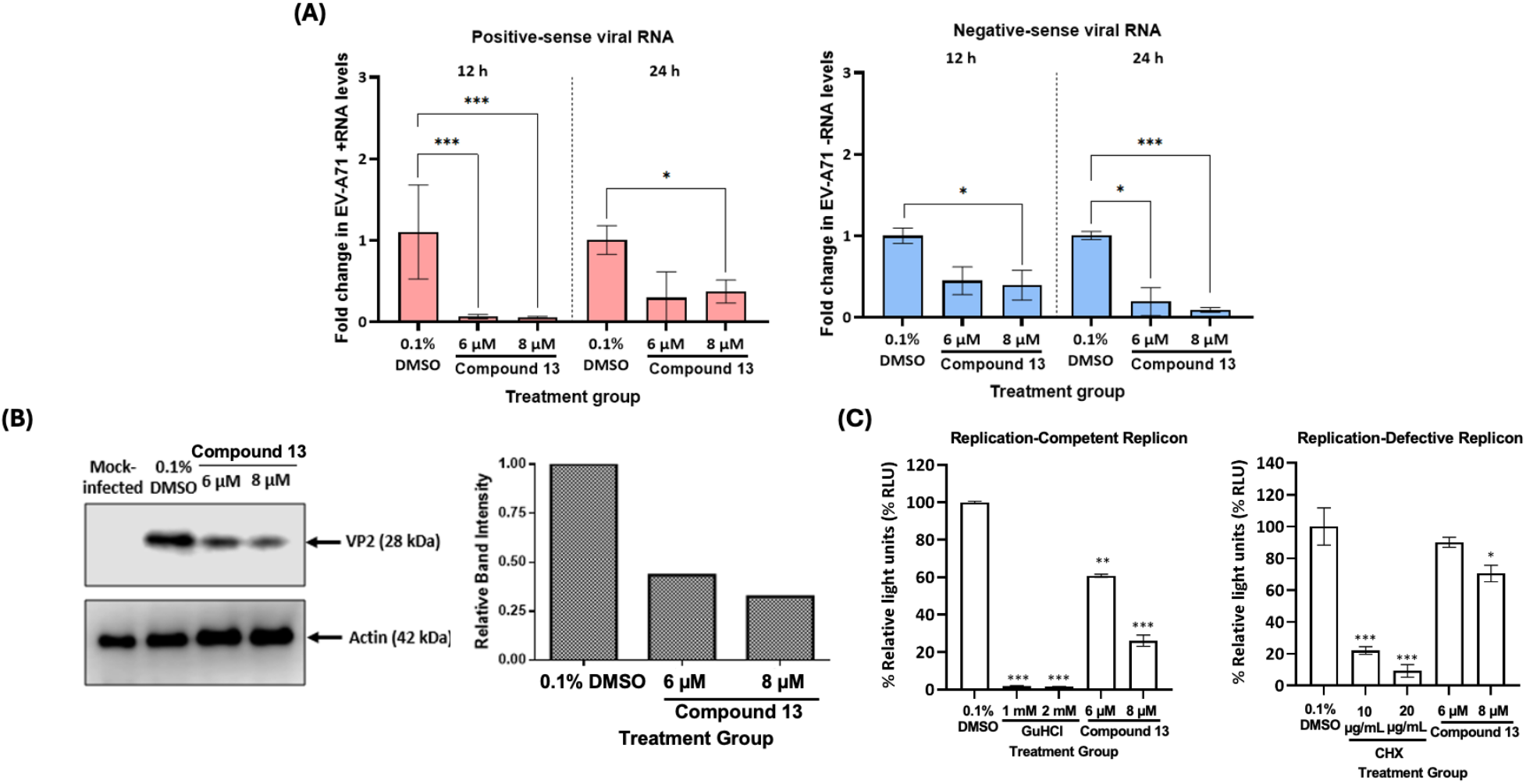
Evaluation of compound **13**’s inhibitory effects against EV-A71 RNA transcription and translation. Lysates of EV-A71 (MOI 1)-infected SH-SY5Y cells treated with specific concentrations of compound **13** were used for (A) qRT-PCR-based viral strand-specific RNA quantification or (B) immunoblotting measurement of specific protein levels. (C) Replicon-based EV-A71 transcription and translation machinery analyses in the presence or absence of compound **13** treatment. Guanidine hydrochloride (GuHCl), an eukaryotic RNA transcription inhibitor^41^, and cycloheximide (CHX), an inhibitor of the elongation phase during RNA translation^42^, were used as positive controls of respective assays. 0.1% DMSO served as the mock control group of respective assays. Luminescence readings generated by SH-SY5Y cells transfected with either the established replication-competent or replication-defective replicons^27^ were measured using a microplate reader and normalized against the DMSO control. Error bars represent the standard deviation from the mean of triplicates.

## 4. Discussion

Millions of paediatric HFMD casess are reported annually across the globe^35,36^. Although HFMD is considered to be a major public health concern with high prevalence, there is currently no effective U.S. Food and Drug Administration (FDA)-approved antiviral therapy for this disease^37^. This has led to substantial socioeconomic effects, with multi-million-dollar annual economic losses associated with HFMD in addition to negative impacts on the physical and mental well-being of infected children and their surrounding caregivers^35,38,39^. As such, there is an urgent need for the discovery and development of potential therapeutic agents to combat widespread enterovirus infections.

Our primary objective in this study was to assess a library of rocaglate/aglain derivatives in a phenotypic high-throughput screen to identify promising hit compound(s) which could be further evaluated for antiviral activities against EV-A71. After excluding all antiviral rocaglate hits from subsequent investigation due to their more prominent cytotoxicity profiles, we identified aglain **13** as the sole hit from the screening assays **(Figures 2, 4A, & 5A)**. By expanding our hit criteria, we were able to identify four near neighbour aglains with similar, albeit weaker viral inhibition. While the limited scope of close structural neighbours available precludes extensive SAR analysis, we noted two unique structural features of compound **13** to be the bridge (C8a) hydroxyl chirality and the C2 pentafluorophenyl (PFP) ring. There are literature reports of PFP rings in bioactive molecules^30-32^. Notably, we have shown that C8a ketones such as **14, 15**, and **18** (**Figure 4**) can form hydrates, leading to positional mimicry of both hydroxyl epimers^25^.

Our TOA findings **(Figure 6A)** suggest that one or more crucial steps of the EV-A71 replication cycle – virus attachment, entry, or intracellular replication – could potentially be targeted by compound **13**. Hence, we first investigated whether compound **13** interferes with the early events of the virus life cycle *via* pre-treatment and co-treatment assays. In the pre-treatment set-up, host cells were first treated with the compound prior to virus challenge. This would allow us to evaluate the ability of compound **13** to competitively inhibit virus particles from binding to surface receptors on target cells, thereby hampering virus entry. Conversely, in the co-treatment assay extracellular virions were treated with the compound prior to infecting the cells. The objective of this set-up was to evaluate the ability of compound **13** to bind vital EV-A71 structural proteins required for virus attachment with host receptors, which will then block viral entry. Interestingly, no significant inhibition of virus yields were detected from either assay **(Figures 6B & 6C)**, suggesting that compound **13**’s antiviral effects against EV-A71 do not involve the initial infection stage of the virus.

Having excluded the virus attachment and entry steps, we deemed it likely that aglain **13** targets the early post-entry stages of EV-A71 life cycle involving the initial virus uncoating, viral RNA genome replication or protein production. As packaging and secretion of newly synthesized EV-A71 virions from infected cells generally begin after 6 h.p.i.^40^, compound **13** is unlikely to interfere with the later phases of the virus life cycle (**Figure 6A**). The ability of compound **13** to interfere with EV-A71 post-entry events was further corroborated *via* our entry-bypass assay wherein host cells were first transfected with *in vitro* isolated viral RNA prior to compound treatment **(Figure 6D)**. The rationale of this entry-bypass approach was to simulate intracellular viral replication steps following virus entry as EV-A71 genetic content was directly introduced into the cells *via* transfection, hence bypassing the early stages of the virus life cycle. As compound **13** targets the intracellular replication steps of EV-A71, its effects on two primary pre-packaging stages of the virus life cycle, namely viral RNA replication and protein production, were evaluated.

We first investigated the effects of compound **13** treatment towards EV-A71 positive- and negative-strand RNA synthesis. Being a positive-strand RNA virus, the initial step of EV-A71 genomic replication involves the synthesis of complementary negative-strand RNA using its positive-strand molecule as the template, followed by rounds of exponential replication to generate new viral RNA components. It was observed that compound **13** was an effective inhibitor of EV-A71 RNA transcription and replication **(Figure 7A)**. The compound’s anti-EV-A71 effects were also reflected by the reduced expression levels of a specific viral protein following treatment **(Figure 7B)**. This could either be due to the compound’s direct inhibition on viral translation machinery, or indirect inhibition *via* viral RNA suppression.

Given the intrinsic association between viral RNA transcription and translation, we utilized our previously established EV-A71 replication-competent or replication-defective replicon constructs which enable a clear distinction of a specific compound’s inhibitory effects against either step of virus replication^27^. Briefly, the replication-competent RNA replicon harbours an intact coding region for EV-A71 3D polymerase, in which 159 nucleotides are deleted within the replication-defective construct. As a result, following an IRES-driven translation in transfected cells, the replication-competent replicon will express luciferase signals generated from both the original replicons as well as newly replicated copies of the RNA. In contrast, impairment of RNA replication capacity in the replication-defective replicon will only result in detection of luciferase signals generated from the original construct, providing specific indication of the compound’s downregulating effects on viral RNA translation. Our replicon data suggested that compound **13**’s antiviral activity against EV-A71 extends beyond the respective individual stages of the virus replication, affecting both RNA transcription and translation **(Figure 7C)**. In fact, stronger suppression of luciferase signals were observed from the replication-competent replicon **(Figure 7C)**, suggesting that interference of viral RNA transcription may play a major role in the compound’s antiviral effects. This may indicate the involvement of one or more unexplored mechanism(s) of action which contribute to the inhibitory efficacy of aglain **13** against EV-A71 replication, contrasting the established rocaglate-associated cellular pathways involving eIF4a^9^ and prohibitin^11^.

In conclusion, findings presented in this study warrant an in-depth mechanistic investigation of the underlying pathway(s) involved in compound **13**’s antiviral activity against major human RNA viruses other than EV-A71. Further evaluation in relevant animal models and medicinal chemistry optimization of both potency and pharmacological properties are necessary prior to potential clinical development of this novel synthetic aglain chemotype.

## Data availability statement

The data that support the findings of this study are available from the corresponding author upon reasonable request.

## Author Contributions

**Conceptualization**: Adrian Oo, Lauren E. Brown, John A. Porco, Jr., Justin Jang Hann Chu. **Data curation**: Adrian Oo, Angel Borge, Regina Ching Hua Lee, Cyrill Kafi Salim, Wenyu Wang, Michael Ricca, Deborah Yuhui Fong. **Methodology**: Adrian Oo, Michael Ricca, Lauren E. Brown, John A. Porco, Jr., Justin Jang Hann Chu. **Investigation**: Adrian Oo, Angel Borge, Regina Ching Hua Lee, Cyrill Kafi Salim, Wenyu Wang, Michael Ricca, Deborah Yuhui Fong. **Project Administration**: Adrian Oo, Lauren E. Brown, John A. Porco, Jr., Justin Jang Hann Chu. **Supervision**: Lauren E. Brown, John A. Porco, Jr., Justin Jang Hann Chu. **Resources**: Sylvie Alonso, Lauren E. Brown, John A. Porco, Jr., Justin Jang Hann Chu. **Funding acquisition**: Sylvie Alonso, Lauren E. Brown, John A. Porco, Jr., Justin Jang Hann Chu. **Writing – original draft**: Adrian Oo, Angel Borge, Michael Ricca, Lauren E. Brown, John A Porco, Jr., Justin Jang Hann Chu. **Writing - review & editing**: Adrian Oo, Sylvie Alonso, Lauren E. Brown, John A Porco, Jr., Justin Jang Hann Chu. All authors read and commented on the final manuscript.

## Acknowledgements

This work was supported by Singapore National Research Foundation Competitive Research Programme (NRF-CRP21-2018-0004) (SA & JJHC), Singapore Ministry of Education (MOE) Academic Research Fund (AcRF) Tier 2 (MOE-T2EP30221-0005) (JJHC), NIH grants R35GM118173 and U01TR002625 (JAP, Jr.). We thank the National Science Foundation for support of NMR (CHE-0619339) and MS (CHE-0443618) facilities at BU and the NIH (S10OD028585) for support of the single-crystal XRD system.

## Conflict of interest disclosure

The authors declare the following competing financial interest(s): J.J. H.C., A.O., W.W., L.E.B., and J.A.P., Jr. are named as inventors on a U.S. provisional patent application pertaining to the findings reported here.

