## Supplementary Information for "*In vitro* discovery of a therapeutic lead for HFMD from a library screen of rocaglates/aglains"

### Table of Contents:

I. Single-crystal X-ray diffraction data for aglain **13** .......... 2

II. Validation of cytotoxicity and antiviral profiles of compound **13** .......... 12

**X-ray crystallographic data for aglain 13**

Crystals of aglain **13** suitable for X-ray analysis were obtained by slow evaporation of methanol. Crystallographic data have been deposited with the Cambridge Crystallograhic Data Centre (CCDC 2383091). Copies of the data can be obtained free of charge on application to the CCDC, 12 Union Road, Cambridge CB21EZ, UK (fax: (+44)-1223-336-033;).

**Computing details**

Data collection : APEX4, (Bruker, 2016); Cell refinement: *CrysAlis PRO* 1.171.42.49 (Rigaku OD, 2022); data reduction: *CrysAlis PRO* 1.171.42.49 (Rigaku OD, 2022); program(s) used to solve structure: SHELXT (Sheldrick, 2015); program(s) used to refine structure: *SHELXL* 2018/3 (Sheldrick, 2015); molecular graphics: Olex2 1.5 (Dolomanov *et al.*, 2009); software used to prepare material for publication: Mercury.

(aglain **13)**

*Crystal data*

| | |
|---|---|
| $C_{32}H_{25}F_5O_6 \cdot CH_4O$·[+solvent] | $Z = 2$ |
| $M_r = 632.56$ | $F(000) = 656$ |
| Triclinic, $P\bar{1}$ | $D_x = 1.273$ Mg m$^{-3}$ |
| $a = 11.2346$ (2) Å | Cu $K\alpha$ radiation, $\lambda = 1.54184$ Å |
| $b = 12.3867$ (3) Å | Cell parameters from 56082 reflections |
| $c = 12.5593$ (3) Å | $\theta = 3.7$–$75.7°$ |
| $\alpha = 105.461$ (2)° | $\mu = 0.92$ mm$^{-1}$ |
| $\beta = 95.6551$ (18)° | $T = 100$ K |
| $\gamma = 98.1647$ (19)° | Block, clear colourless |
| $V = 1650.30$ (7) Å$^3$ | 0.2 × 0.2 × 0.2 mm |

*Data collection*

| | |
|---|---|
| Bruker D8 Venture diffractometer | 5797 reflections with $I > 2\sigma(I)$ |
| Radiation source: microfocus sealed tube | $R_{int} = 0.119$ |
| ω and ϕ scans | $\theta_{max} = 68.2°$, $\theta_{min} = 3.7°$ |
| Absorption correction: multi-scan *CrysAlis PRO* 1.171.42.49 (Rigaku Oxford Diffraction, 2022) Empirical absorption | $h = -13 \rightarrow 13$ |

| | |
|---|---|
| correction using spherical harmonics, implemented in SCALE3 ABSPACK scaling algorithm. | |
| $T_{min}$ = 0.735, $T_{max}$ = 1.000 | $k$ = -14→14 |
| 68999 measured reflections | $l$ = -14→14 |
| 5878 independent reflections | |

*Refinement*

| | |
|---|---|
| Refinement on $F^2$ | Hydrogen site location: mixed |
| Least-squares matrix: full | H atoms treated by a mixture of independent and constrained refinement |
| $R[F^2 > 2\sigma(F^2)]$ = 0.065 | $w = 1/[\sigma^2(F_o^2) + (0.0942P)^2 + 1.369P]$ where $P = (F_o^2 + 2F_c^2)/3$ |
| $wR(F^2)$ = 0.176 | $(\Delta/\sigma)_{max}$ < 0.001 |
| $S$ = 1.11 | $\Delta\rangle_{max}$ = 0.40 e Å$^{-3}$ |
| 5878 reflections | $\Delta\rangle_{min}$ = -0.43 e Å$^{-3}$ |
| 423 parameters | Extinction correction: *SHELXL2018*/3 (Sheldrick 2018), $Fc^*=kFc[1+0.001xFc^2\lambda^3/\sin(2\theta)]^{-1/4}$ |
| 0 restraints | Extinction coefficient: 0.0146 (14) |
| Primary atom site location: structure-invariant direct methods | |

*Special details*

| |
|---|
| *Geometry*. All esds (except the esd in the dihedral angle between two l.s. planes) are estimated using the full covariance matrix. The cell esds are taken into account individually in the estimation of esds in distances, angles and torsion angles; correlations between esds in cell parameters are only used when they are defined by crystal symmetry. An approximate (isotropic) treatment of cell esds is used for estimating esds involving l.s. planes. |

*Fractional atomic coordinates and isotropic or equivalent isotropic displacement parameters (Å$^2$) for aglain **13***

| | $x$ | $y$ | $z$ | $U_{iso}$*/$U_{eq}$ |
|---|---|---|---|---|
| F5 | 0.14572 (11) | 0.54539 (12) | 0.62057 (11) | 0.0310 (3) |
| F1 | 0.10383 (12) | 0.35724 (11) | 0.90201 (11) | 0.0324 (3) |
| F4 | -0.06934 (12) | 0.60452 (13) | 0.64542 (12) | 0.0402 (4) |
| O3 | 0.50547 (12) | 0.56387 (11) | 0.69210 (11) | 0.0190 (3) |
| F2 | -0.11289 (12) | 0.41428 (13) | 0.92153 (11) | 0.0399 (4) |

| | | | | |
|---|---|---|---|---|
| O5 | 0.57617 (13) | 0.56783 (13) | 0.90579 (12) | 0.0232 (3) |
| O4 | 0.32997 (13) | 0.55440 (13) | 0.96748 (12) | 0.0235 (3) |
| F3 | -0.19982 (12) | 0.54538 (14) | 0.79952 (12) | 0.0440 (4) |
| O1 | 0.39807 (14) | 0.92125 (13) | 0.69217 (13) | 0.0292 (4) |
| O6 | 0.84524 (14) | 0.20665 (13) | 0.58730 (14) | 0.0315 (4) |
| O2 | 0.21809 (14) | 0.71294 (13) | 0.92070 (13) | 0.0291 (4) |
| O7 | 0.5957 (2) | 0.76842 (17) | 1.07864 (16) | 0.0468 (5) |
| C19 | 0.29792 (17) | 0.28884 (17) | 0.59230 (17) | 0.0211 (4) |
| C12 | 0.56461 (17) | 0.38498 (17) | 0.67626 (16) | 0.0198 (4) |
| C3 | 0.44281 (17) | 0.64859 (16) | 0.73657 (15) | 0.0188 (4) |
| C2 | 0.46097 (18) | 0.74302 (17) | 0.69621 (16) | 0.0206 (4) |
| H2 | 0.518592 | 0.749484 | 0.646744 | 0.025* |
| C13 | 0.62293 (19) | 0.39914 (17) | 0.58671 (17) | 0.0221 (4) |
| H13 | 0.599034 | 0.450402 | 0.547768 | 0.027* |
| C9 | 0.35472 (18) | 0.53420 (17) | 0.85573 (16) | 0.0202 (4) |
| C26 | 0.26724 (18) | 0.43191 (17) | 0.77236 (16) | 0.0201 (4) |
| H26 | 0.269027 | 0.365272 | 0.802847 | 0.024* |
| C25 | 0.33340 (17) | 0.40910 (17) | 0.66780 (16) | 0.0197 (4) |
| H25 | 0.307932 | 0.460118 | 0.623399 | 0.024* |
| C4 | 0.36491 (18) | 0.63939 (17) | 0.81468 (16) | 0.0201 (4) |
| C11 | 0.47231 (17) | 0.45714 (16) | 0.71857 (16) | 0.0185 (4) |
| C1 | 0.39248 (19) | 0.82744 (17) | 0.73029 (17) | 0.0235 (4) |
| C17 | 0.5982 (2) | 0.30512 (18) | 0.72821 (18) | 0.0248 (4) |
| H17 | 0.557313 | 0.291291 | 0.787232 | 0.030* |
| C10 | 0.47291 (18) | 0.48795 (17) | 0.84570 (16) | 0.0199 (4) |
| H10 | 0.467633 | 0.417533 | 0.870851 | 0.024* |
| C32 | 0.08689 (19) | 0.51212 (18) | 0.69782 (17) | 0.0245 (4) |
| C27 | 0.13756 (18) | 0.45112 (17) | 0.76272 (17) | 0.0221 (4) |
| C20 | 0.25911 (19) | 0.27143 (18) | 0.47958 (18) | 0.0249 (4) |
| H20 | 0.261185 | 0.335143 | 0.450802 | 0.030* |
| C6 | 0.3096 (2) | 0.82031 (19) | 0.80509 (18) | 0.0265 (5) |
| H6 | 0.262513 | 0.878294 | 0.826965 | 0.032* |
| C24 | 0.2963 (2) | 0.19369 (18) | 0.63203 (19) | 0.0273 (5) |
| H24 | 0.323424 | 0.203293 | 0.708582 | 0.033* |
| C23 | 0.2551 (2) | 0.08471 (19) | 0.5602 (2) | 0.0314 (5) |
| H23 | 0.254999 | 0.020630 | 0.588224 | 0.038* |
| C14 | 0.71544 (19) | 0.33975 (18) | 0.55315 (17) | 0.0236 (4) |

| | | | | |
|---|---|---|---|---|
| H14 | 0.754069 | 0.350755 | 0.491948 | 0.028* |
| C15 | 0.75118 (19) | 0.26461 (17) | 0.60901 (18) | 0.0236 (4) |
| C31 | -0.02664 (19) | 0.5432 (2) | 0.70878 (19) | 0.0290 (5) |
| C21 | 0.2173 (2) | 0.1628 (2) | 0.40802 (19) | 0.0315 (5) |
| H21 | 0.190749 | 0.152757 | 0.331268 | 0.038* |
| C28 | 0.06423 (19) | 0.41827 (18) | 0.83610 (17) | 0.0248 (4) |
| C5 | 0.29738 (19) | 0.72722 (18) | 0.84669 (17) | 0.0239 (4) |
| C29 | -0.0484 (2) | 0.4477 (2) | 0.84749 (18) | 0.0298 (5) |
| C7 | 0.4931 (2) | 0.9404 (2) | 0.6276 (2) | 0.0332 (5) |
| H7A | 0.477984 | 0.880808 | 0.556106 | 0.050* |
| H7B | 0.571298 | 0.938574 | 0.668525 | 0.050* |
| H7C | 0.494811 | 1.014856 | 0.613940 | 0.050* |
| C30 | -0.09357 (19) | 0.5123 (2) | 0.78565 (19) | 0.0313 (5) |
| C22 | 0.2145 (2) | 0.06880 (19) | 0.4491 (2) | 0.0329 (5) |
| H22 | 0.184914 | -0.005648 | 0.400970 | 0.039* |
| C16 | 0.6899 (2) | 0.24581 (18) | 0.69528 (19) | 0.0275 (5) |
| H16 | 0.711339 | 0.191821 | 0.731772 | 0.033* |
| C18 | 0.9103 (2) | 0.2233 (2) | 0.4999 (2) | 0.0351 (5) |
| H18A | 0.855642 | 0.196513 | 0.428855 | 0.053* |
| H18B | 0.977507 | 0.180511 | 0.495871 | 0.053* |
| H18C | 0.942741 | 0.304477 | 0.514769 | 0.053* |
| C8 | 0.1471 (3) | 0.8005 (3) | 0.9573 (3) | 0.0539 (8) |
| H8A | 0.096526 | 0.808496 | 0.892587 | 0.081* |
| H8B | 0.201484 | 0.872799 | 0.994739 | 0.081* |
| H8C | 0.094903 | 0.779723 | 1.009379 | 0.081* |
| C33 | 0.5381 (5) | 0.8561 (3) | 1.0656 (3) | 0.0858 (15) |
| H33A | 0.554425 | 0.872810 | 0.995872 | 0.129* |
| H33B | 0.568554 | 0.923977 | 1.128584 | 0.129* |
| H33C | 0.450364 | 0.833852 | 1.062989 | 0.129* |
| H7 | 0.575 (3) | 0.715 (3) | 1.025 (3) | 0.049 (9)* |
| H5 | 0.617 (3) | 0.536 (3) | 0.943 (3) | 0.050 (9)* |
| H4 | 0.275 (4) | 0.596 (3) | 0.986 (3) | 0.060 (10)* |

*Atomic displacement parameters (Å$^2$) for aglain **13***

| | $U^{11}$ | $U^{22}$ | $U^{33}$ | $U^{12}$ | $U^{13}$ | $U^{23}$ |
|---|---|---|---|---|---|---|
| F5 | 0.0235 (6) | 0.0471 (8) | 0.0339 (7) | 0.0112 (5) | 0.0098 (5) | 0.0262 (6) |

| | | | | | | |
|---|---|---|---|---|---|---|
| F1 | 0.0329 (7) | 0.0397 (7) | 0.0319 (7) | 0.0039 (6) | 0.0125 (5) | 0.0209 (6) |
| F4 | 0.0266 (7) | 0.0589 (9) | 0.0435 (8) | 0.0179 (6) | 0.0030 (6) | 0.0240 (7) |
| O3 | 0.0202 (7) | 0.0204 (7) | 0.0205 (7) | 0.0059 (5) | 0.0081 (5) | 0.0099 (5) |
| F2 | 0.0260 (7) | 0.0608 (9) | 0.0324 (7) | -0.0019 (6) | 0.0164 (6) | 0.0133 (7) |
| O5 | 0.0197 (7) | 0.0313 (8) | 0.0207 (7) | 0.0044 (6) | 0.0024 (5) | 0.0112 (6) |
| O4 | 0.0244 (8) | 0.0331 (8) | 0.0188 (7) | 0.0091 (6) | 0.0095 (6) | 0.0126 (6) |
| F3 | 0.0189 (7) | 0.0665 (10) | 0.0429 (8) | 0.0114 (6) | 0.0073 (6) | 0.0064 (7) |
| O1 | 0.0345 (8) | 0.0283 (8) | 0.0356 (9) | 0.0129 (6) | 0.0151 (7) | 0.0198 (7) |
| O6 | 0.0277 (8) | 0.0315 (8) | 0.0416 (9) | 0.0131 (7) | 0.0148 (7) | 0.0133 (7) |
| O2 | 0.0304 (8) | 0.0326 (8) | 0.0355 (8) | 0.0159 (7) | 0.0213 (7) | 0.0172 (7) |
| O7 | 0.0691 (14) | 0.0383 (10) | 0.0292 (9) | 0.0038 (9) | 0.0093 (9) | 0.0051 (8) |
| C19 | 0.0157 (9) | 0.0263 (10) | 0.0252 (10) | 0.0065 (8) | 0.0068 (7) | 0.0112 (8) |
| C12 | 0.0173 (9) | 0.0225 (10) | 0.0204 (9) | 0.0021 (7) | 0.0033 (7) | 0.0081 (7) |
| C3 | 0.0172 (9) | 0.0230 (10) | 0.0168 (9) | 0.0043 (7) | 0.0020 (7) | 0.0064 (7) |
| C2 | 0.0206 (10) | 0.0246 (10) | 0.0188 (9) | 0.0041 (8) | 0.0059 (7) | 0.0088 (8) |
| C13 | 0.0233 (10) | 0.0232 (10) | 0.0214 (10) | 0.0041 (8) | 0.0043 (8) | 0.0084 (8) |
| C9 | 0.0197 (10) | 0.0276 (10) | 0.0171 (9) | 0.0052 (8) | 0.0063 (7) | 0.0110 (8) |
| C26 | 0.0197 (10) | 0.0239 (10) | 0.0213 (10) | 0.0045 (8) | 0.0064 (7) | 0.0125 (8) |
| C25 | 0.0186 (9) | 0.0238 (10) | 0.0217 (10) | 0.0059 (7) | 0.0063 (7) | 0.0128 (8) |
| C4 | 0.0194 (9) | 0.0235 (10) | 0.0190 (9) | 0.0038 (8) | 0.0040 (7) | 0.0083 (8) |
| C11 | 0.0192 (9) | 0.0226 (10) | 0.0177 (9) | 0.0039 (7) | 0.0054 (7) | 0.0115 (7) |
| C1 | 0.0257 (10) | 0.0226 (10) | 0.0240 (10) | 0.0041 (8) | 0.0028 (8) | 0.0102 (8) |
| C17 | 0.0271 (11) | 0.0280 (11) | 0.0256 (10) | 0.0073 (8) | 0.0102 (8) | 0.0146 (8) |
| C10 | 0.0190 (10) | 0.0251 (10) | 0.0187 (9) | 0.0052 (8) | 0.0056 (7) | 0.0096 (8) |
| C32 | 0.0204 (10) | 0.0322 (11) | 0.0227 (10) | 0.0028 (8) | 0.0063 (8) | 0.0106 (8) |
| C27 | 0.0187 (10) | 0.0260 (10) | 0.0221 (10) | 0.0018 (8) | 0.0045 (8) | 0.0080 (8) |
| C20 | 0.0235 (10) | 0.0280 (11) | 0.0272 (11) | 0.0075 (8) | 0.0065 (8) | 0.0119 (8) |
| C6 | 0.0273 (11) | 0.0284 (11) | 0.0280 (11) | 0.0102 (9) | 0.0076 (8) | 0.0110 (9) |
| C24 | 0.0259 (11) | 0.0294 (11) | 0.0305 (11) | 0.0052 (9) | 0.0046 (8) | 0.0148 (9) |
| C23 | 0.0299 (12) | 0.0238 (11) | 0.0452 (14) | 0.0067 (9) | 0.0103 (10) | 0.0150 (10) |
| C14 | 0.0236 (10) | 0.0258 (10) | 0.0223 (10) | 0.0027 (8) | 0.0093 (8) | 0.0070 (8) |
| C15 | 0.0204 (10) | 0.0226 (10) | 0.0281 (11) | 0.0054 (8) | 0.0062 (8) | 0.0058 (8) |
| C31 | 0.0206 (11) | 0.0378 (12) | 0.0288 (11) | 0.0064 (9) | 0.0003 (8) | 0.0105 (9) |
| C21 | 0.0319 (12) | 0.0357 (12) | 0.0251 (11) | 0.0071 (9) | 0.0036 (9) | 0.0051 (9) |
| C28 | 0.0228 (10) | 0.0293 (11) | 0.0219 (10) | -0.0010 (8) | 0.0059 (8) | 0.0083 (8) |
| C5 | 0.0217 (10) | 0.0310 (11) | 0.0227 (10) | 0.0073 (8) | 0.0085 (8) | 0.0105 (8) |
| C29 | 0.0228 (11) | 0.0400 (12) | 0.0225 (10) | -0.0049 (9) | 0.0074 (8) | 0.0055 (9) |

| | | | | | | |
|---|---|---|---|---|---|---|
| C7 | 0.0370 (13) | 0.0298 (11) | 0.0432 (13) | 0.0081 (10) | 0.0164 (10) | 0.0229 (10) |
| C30 | 0.0149 (10) | 0.0435 (13) | 0.0300 (11) | 0.0033 (9) | 0.0042 (8) | 0.0017 (10) |
| C22 | 0.0295 (12) | 0.0246 (11) | 0.0427 (13) | 0.0058 (9) | 0.0087 (10) | 0.0047 (9) |
| C16 | 0.0283 (11) | 0.0272 (11) | 0.0323 (11) | 0.0090 (9) | 0.0074 (9) | 0.0144 (9) |
| C18 | 0.0311 (12) | 0.0354 (12) | 0.0426 (13) | 0.0106 (10) | 0.0196 (10) | 0.0102 (10) |
| C8 | 0.0642 (19) | 0.0514 (16) | 0.074 (2) | 0.0378 (15) | 0.0554 (17) | 0.0350 (15) |
| C33 | 0.174 (5) | 0.0380 (17) | 0.0413 (18) | 0.033 (2) | -0.008 (2) | 0.0051 (13) |

*Geometric parameters (Å, º) for aglain **13***

| | | | |
|---|---|---|---|
| F5—C32 | 1.345 (2) | C4—C5 | 1.408 (3) |
| F1—C28 | 1.346 (3) | C11—C10 | 1.539 (3) |
| F4—C31 | 1.338 (3) | C1—C6 | 1.397 (3) |
| O3—C3 | 1.372 (2) | C17—H17 | 0.9500 |
| O3—C11 | 1.454 (2) | C17—C16 | 1.382 (3) |
| F2—C29 | 1.347 (3) | C10—H10 | 1.0000 |
| O5—C10 | 1.413 (2) | C32—C27 | 1.385 (3) |
| O5—H5 | 0.83 (4) | C32—C31 | 1.393 (3) |
| O4—C9 | 1.421 (2) | C27—C28 | 1.396 (3) |
| O4—H4 | 0.87 (4) | C20—H20 | 0.9500 |
| F3—C30 | 1.329 (3) | C20—C21 | 1.390 (3) |
| O1—C1 | 1.366 (2) | C6—H6 | 0.9500 |
| O1—C7 | 1.433 (3) | C6—C5 | 1.383 (3) |
| O6—C15 | 1.369 (3) | C24—H24 | 0.9500 |
| O6—C18 | 1.420 (3) | C24—C23 | 1.393 (3) |
| O2—C5 | 1.377 (2) | C23—H23 | 0.9500 |
| O2—C8 | 1.441 (3) | C23—C22 | 1.377 (4) |
| O7—C33 | 1.378 (4) | C14—H14 | 0.9500 |
| O7—H7 | 0.79 (4) | C14—C15 | 1.385 (3) |
| C19—C25 | 1.510 (3) | C15—C16 | 1.392 (3) |
| C19—C20 | 1.389 (3) | C31—C30 | 1.379 (3) |
| C19—C24 | 1.396 (3) | C21—H21 | 0.9500 |
| C12—C13 | 1.394 (3) | C21—C22 | 1.392 (3) |
| C12—C11 | 1.508 (3) | C28—C29 | 1.376 (3) |
| C12—C17 | 1.396 (3) | C29—C30 | 1.367 (4) |
| C3—C2 | 1.391 (3) | C7—H7A | 0.9800 |
| C3—C4 | 1.393 (3) | C7—H7B | 0.9800 |

| | | | |
|---|---|---|---|
| C2—H2 | 0.9500 | C7—H7C | 0.9800 |
| C2—C1 | 1.387 (3) | C22—H22 | 0.9500 |
| C13—H13 | 0.9500 | C16—H16 | 0.9500 |
| C13—C14 | 1.392 (3) | C18—H18A | 0.9800 |
| C9—C26 | 1.550 (3) | C18—H18B | 0.9800 |
| C9—C4 | 1.518 (3) | C18—H18C | 0.9800 |
| C9—C10 | 1.522 (3) | C8—H8A | 0.9800 |
| C26—H26 | 1.0000 | C8—H8B | 0.9800 |
| C26—C25 | 1.553 (3) | C8—H8C | 0.9800 |
| C26—C27 | 1.508 (3) | C33—H33A | 0.9800 |
| C25—H25 | 1.0000 | C33—H33B | 0.9800 |
| C25—C11 | 1.592 (3) | C33—H33C | 0.9800 |
| C3—O3—C11 | 115.80 (14) | C19—C20—H20 | 119.3 |
| C10—O5—H5 | 108 (2) | C19—C20—C21 | 121.4 (2) |
| C9—O4—H4 | 117 (2) | C21—C20—H20 | 119.3 |
| C1—O1—C7 | 117.03 (16) | C1—C6—H6 | 120.6 |
| C15—O6—C18 | 117.77 (17) | C5—C6—C1 | 118.72 (19) |
| C5—O2—C8 | 117.94 (17) | C5—C6—H6 | 120.6 |
| C33—O7—H7 | 109 (2) | C19—C24—H24 | 119.8 |
| C20—C19—C25 | 119.09 (18) | C23—C24—C19 | 120.5 (2) |
| C20—C19—C24 | 118.16 (19) | C23—C24—H24 | 119.8 |
| C24—C19—C25 | 122.66 (18) | C24—C23—H23 | 119.6 |
| C13—C12—C11 | 121.02 (17) | C22—C23—C24 | 120.8 (2) |
| C13—C12—C17 | 117.71 (19) | C22—C23—H23 | 119.6 |
| C17—C12—C11 | 121.22 (17) | C13—C14—H14 | 120.0 |
| O3—C3—C2 | 115.06 (17) | C15—C14—C13 | 119.99 (19) |
| O3—C3—C4 | 122.30 (17) | C15—C14—H14 | 120.0 |
| C2—C3—C4 | 122.63 (18) | O6—C15—C14 | 124.95 (19) |
| C3—C2—H2 | 121.0 | O6—C15—C16 | 115.86 (18) |
| C1—C2—C3 | 118.06 (18) | C14—C15—C16 | 119.18 (19) |
| C1—C2—H2 | 121.0 | F4—C31—C32 | 119.5 (2) |
| C12—C13—H13 | 119.3 | F4—C31—C30 | 120.4 (2) |
| C14—C13—C12 | 121.37 (19) | C30—C31—C32 | 120.1 (2) |
| C14—C13—H13 | 119.3 | C20—C21—H21 | 120.1 |
| O4—C9—C26 | 113.46 (16) | C20—C21—C22 | 119.8 (2) |
| O4—C9—C4 | 113.65 (16) | C22—C21—H21 | 120.1 |

| | | | |
|---|---|---|---|
| O4—C9—C10 | 109.53 (15) | F1—C28—C27 | 120.16 (19) |
| C4—C9—C26 | 111.69 (15) | F1—C28—C29 | 116.77 (19) |
| C4—C9—C10 | 108.47 (16) | C29—C28—C27 | 123.1 (2) |
| C10—C9—C26 | 98.92 (15) | O2—C5—C4 | 115.35 (18) |
| C9—C26—H26 | 106.9 | O2—C5—C6 | 122.81 (19) |
| C9—C26—C25 | 102.57 (15) | C6—C5—C4 | 121.83 (19) |
| C25—C26—H26 | 106.9 | F2—C29—C28 | 119.5 (2) |
| C27—C26—C9 | 112.42 (16) | F2—C29—C30 | 120.2 (2) |
| C27—C26—H26 | 106.9 | C30—C29—C28 | 120.3 (2) |
| C27—C26—C25 | 120.28 (16) | O1—C7—H7A | 109.5 |
| C19—C25—C26 | 113.52 (16) | O1—C7—H7B | 109.5 |
| C19—C25—H25 | 106.7 | O1—C7—H7C | 109.5 |
| C19—C25—C11 | 119.26 (16) | H7A—C7—H7B | 109.5 |
| C26—C25—H25 | 106.7 | H7A—C7—H7C | 109.5 |
| C26—C25—C11 | 103.24 (15) | H7B—C7—H7C | 109.5 |
| C11—C25—H25 | 106.7 | F3—C30—C31 | 120.4 (2) |
| C3—C4—C9 | 118.30 (17) | F3—C30—C29 | 120.8 (2) |
| C3—C4—C5 | 117.08 (18) | C29—C30—C31 | 118.9 (2) |
| C5—C4—C9 | 124.58 (18) | C23—C22—C21 | 119.4 (2) |
| O3—C11—C12 | 106.00 (14) | C23—C22—H22 | 120.3 |
| O3—C11—C25 | 107.27 (14) | C21—C22—H22 | 120.3 |
| O3—C11—C10 | 106.42 (15) | C17—C16—C15 | 120.47 (19) |
| C12—C11—C25 | 117.56 (16) | C17—C16—H16 | 119.8 |
| C12—C11—C10 | 114.60 (16) | C15—C16—H16 | 119.8 |
| C10—C11—C25 | 104.33 (15) | O6—C18—H18A | 109.5 |
| O1—C1—C2 | 123.36 (19) | O6—C18—H18B | 109.5 |
| O1—C1—C6 | 115.08 (18) | O6—C18—H18C | 109.5 |
| C2—C1—C6 | 121.55 (19) | H18A—C18—H18B | 109.5 |
| C12—C17—H17 | 119.4 | H18A—C18—H18C | 109.5 |
| C16—C17—C12 | 121.16 (19) | H18B—C18—H18C | 109.5 |
| C16—C17—H17 | 119.4 | O2—C8—H8A | 109.5 |
| O5—C10—C9 | 113.28 (16) | O2—C8—H8B | 109.5 |
| O5—C10—C11 | 112.88 (16) | O2—C8—H8C | 109.5 |
| O5—C10—H10 | 109.9 | H8A—C8—H8B | 109.5 |
| C9—C10—C11 | 100.69 (15) | H8A—C8—H8C | 109.5 |
| C9—C10—H10 | 109.9 | H8B—C8—H8C | 109.5 |
| C11—C10—H10 | 109.9 | O7—C33—H33A | 109.5 |

| | | | |
|---|---|---|---|
| F5—C32—C27 | 121.44 (18) | O7—C33—H33B | 109.5 |
| F5—C32—C31 | 116.15 (19) | O7—C33—H33C | 109.5 |
| C27—C32—C31 | 122.4 (2) | H33A—C33—H33B | 109.5 |
| C32—C27—C26 | 126.28 (18) | H33A—C33—H33C | 109.5 |
| C32—C27—C28 | 115.19 (19) | H33B—C33—H33C | 109.5 |
| C28—C27—C26 | 117.95 (18) | | |
| F5—C32—C27—C26 | 11.5 (3) | C26—C25—C11—C12 | 134.99 (16) |
| F5—C32—C27—C28 | -177.45 (18) | C26—C25—C11—C10 | 6.83 (19) |
| F5—C32—C31—F4 | -1.2 (3) | C26—C27—C28—F1 | -9.1 (3) |
| F5—C32—C31—C30 | 179.76 (19) | C26—C27—C28—C29 | 169.2 (2) |
| F1—C28—C29—F2 | -0.5 (3) | C25—C19—C20—C21 | -175.41 (19) |
| F1—C28—C29—C30 | 178.24 (19) | C25—C19—C24—C23 | 175.69 (19) |
| F4—C31—C30—F3 | -1.6 (3) | C25—C26—C27—C32 | -39.1 (3) |
| F4—C31—C30—C29 | 178.7 (2) | C25—C26—C27—C28 | 150.07 (19) |
| O3—C3—C2—C1 | 174.53 (17) | C25—C11—C10—O5 | -158.59 (15) |
| O3—C3—C4—C9 | 3.2 (3) | C25—C11—C10—C9 | -37.51 (18) |
| O3—C3—C4—C5 | -174.43 (17) | C4—C3—C2—C1 | -4.2 (3) |
| O3—C11—C10—O5 | -45.3 (2) | C4—C9—C26—C25 | 64.49 (19) |
| O3—C11—C10—C9 | 75.73 (17) | C4—C9—C26—C27 | -66.1 (2) |
| F2—C29—C30—F3 | 1.7 (3) | C4—C9—C10—O5 | 57.8 (2) |
| F2—C29—C30—C31 | -178.61 (19) | C4—C9—C10—C11 | -62.96 (18) |
| O4—C9—C26—C25 | -165.47 (15) | C11—O3—C3—C2 | -169.97 (16) |
| O4—C9—C26—C27 | 63.9 (2) | C11—O3—C3—C4 | 8.8 (2) |
| O4—C9—C4—C3 | 148.86 (17) | C11—C12—C13—C14 | 174.52 (18) |
| O4—C9—C4—C5 | -33.7 (3) | C11—C12—C17—C16 | -174.6 (2) |
| O4—C9—C10—O5 | -66.7 (2) | C1—C6—C5—O2 | -179.99 (19) |
| O4—C9—C10—C11 | 172.49 (15) | C1—C6—C5—C4 | -0.8 (3) |
| O1—C1—C6—C5 | 179.29 (19) | C17—C12—C13—C14 | -2.9 (3) |
| O6—C15—C16—C17 | 176.2 (2) | C17—C12—C11—O3 | 149.74 (18) |
| C19—C25—C11—O3 | 127.19 (17) | C17—C12—C11—C25 | -90.4 (2) |
| C19—C25—C11—C12 | 8.0 (2) | C17—C12—C11—C10 | 32.7 (3) |
| C19—C25—C11—C10 | -120.17 (18) | C10—C9—C26—C25 | -49.56 (17) |
| C19—C20—C21—C22 | -0.4 (3) | C10—C9—C26—C27 | 179.84 (16) |
| C19—C24—C23—C22 | -0.5 (3) | C10—C9—C4—C3 | 26.8 (2) |
| C12—C13—C14—C15 | 0.1 (3) | C10—C9—C4—C5 | -155.82 (19) |
| C12—C11—C10—O5 | 71.5 (2) | C32—C27—C28—F1 | 179.07 (18) |

| | | | |
|---|---|---|---|
| C12—C11—C10—C9 | -167.46 (16) | C32—C27—C28—C29 | -2.6 (3) |
| C12—C17—C16—C15 | 0.0 (3) | C32—C31—C30—F3 | 177.4 (2) |
| C3—O3—C11—C12 | -171.62 (15) | C32—C31—C30—C29 | -2.3 (3) |
| C3—O3—C11—C25 | 62.00 (19) | C27—C26—C25—C19 | -78.0 (2) |
| C3—O3—C11—C10 | -49.2 (2) | C27—C26—C25—C11 | 151.49 (17) |
| C3—C2—C1—O1 | -176.76 (18) | C27—C32—C31—F4 | 178.39 (19) |
| C3—C2—C1—C6 | 1.6 (3) | C27—C32—C31—C30 | -0.6 (3) |
| C3—C4—C5—O2 | 177.62 (17) | C27—C28—C29—F2 | -178.90 (19) |
| C3—C4—C5—C6 | -1.6 (3) | C27—C28—C29—C30 | -0.1 (3) |
| C2—C3—C4—C9 | -178.17 (18) | C20—C19—C25—C26 | 127.32 (19) |
| C2—C3—C4—C5 | 4.2 (3) | C20—C19—C25—C11 | -110.7 (2) |
| C2—C1—C6—C5 | 0.8 (3) | C20—C19—C24—C23 | -0.9 (3) |
| C13—C12—C11—O3 | -27.6 (2) | C20—C21—C22—C23 | -1.0 (3) |
| C13—C12—C11—C25 | 92.3 (2) | C24—C19—C25—C26 | -49.2 (3) |
| C13—C12—C11—C10 | -144.63 (18) | C24—C19—C25—C11 | 72.8 (2) |
| C13—C12—C17—C16 | 2.8 (3) | C24—C19—C20—C21 | 1.3 (3) |
| C13—C14—C15—O6 | -176.19 (19) | C24—C23—C22—C21 | 1.4 (3) |
| C13—C14—C15—C16 | 2.8 (3) | C14—C15—C16—C17 | -2.9 (3) |
| C9—C26—C25—C19 | 156.39 (16) | C31—C32—C27—C26 | -168.1 (2) |
| C9—C26—C25—C11 | 25.85 (18) | C31—C32—C27—C28 | 3.0 (3) |
| C9—C26—C27—C32 | 81.8 (2) | C28—C29—C30—F3 | -177.0 (2) |
| C9—C26—C27—C28 | -89.0 (2) | C28—C29—C30—C31 | 2.6 (3) |
| C9—C4—C5—O2 | 0.2 (3) | C7—O1—C1—C2 | -9.9 (3) |
| C9—C4—C5—C6 | -179.09 (19) | C7—O1—C1—C6 | 171.7 (2) |
| C26—C9—C4—C3 | -81.2 (2) | C18—O6—C15—C14 | -0.9 (3) |
| C26—C9—C4—C5 | 96.2 (2) | C18—O6—C15—C16 | -179.98 (19) |
| C26—C9—C10—O5 | 174.38 (15) | C8—O2—C5—C4 | 179.5 (2) |
| C26—C9—C10—C11 | 53.59 (17) | C8—O2—C5—C6 | -1.2 (3) |
| C26—C25—C11—O3 | -105.81 (15) | | |

*Hydrogen-bond geometry (Å, º) for aglain* ***13***

| *D*—H···*A* | *D*—H | H···*A* | *D*···*A* | *D*—H···*A* |
|---|---|---|---|---|
| O7—H7···O5 | 0.79 (4) | 2.03 (4) | 2.795 (2) | 161 (3) |
| O5—H5···O4$^{i}$ | 0.83 (4) | 1.90 (4) | 2.710 (2) | 165 (3) |
| O4—H4···O2 | 0.87 (4) | 2.00 (4) | 2.637 (2) | 130 (3) |

Symmetry code: (i) $-x+1$, $-y+1$, $-z+2$.

Document origin: *publCIF* [Westrip, S. P. (2010). *J. Apply. Cryst.*, **43**, 920-925].

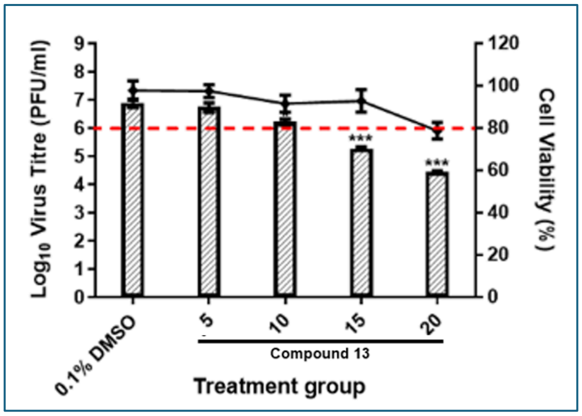

**Figure S1.** Validation of cytotoxicity and antiviral profiles of compound **13**. NSC-34 cells were treated with specific concentrations of compound **13**, in the presence or absence of EV-A71 (MOI 1) infection. Resulting cell viability and virus yield from respective experimental setups were determined via alamarBlue™ and plaque assays, respectively.
